# KLK4 inhibition by cyclic and acyclic peptides: structural and dynamical insights into standard-mechanism protease inhibitors

**DOI:** 10.1101/570390

**Authors:** Blake T. Riley, Olga Ilyichova, Simon J. de Veer, Joakim E. Swedberg, Emily Wilson, David E. Hoke, Jonathan M. Harris, Ashley M. Buckle

**Affiliations:** Department of Biochemistry and Molecular Biology, Biomedicine Discovery Institute, Monash University, Clayton, Victoria 3800, Australia; Institute for Molecular Bioscience, The University of Queensland, Brisbane, QLD 4072, Australia; Institute of Health and Biomedical Innovation, Queensland University of Technology, Brisbane, Queensland 4059, Australia

**Keywords:** Kallikrein-related peptidase 4, KLK4; Sunflower Trypsin Inhibitor, SFTI; Standard-mechanism, Laskowski-mechanism, Conformational dynamics, protease inhibitor

## Abstract

Sunflower Trypsin Inhibitor (SFTI-1) is a 14-amino acid serine protease inhibitor. The dual anti-parallel β-sheet arrangement of SFTI-1 is stabilized by a N-terminal-C-terminal backbone cyclization and a further disulfide bridge to form a final bicyclic structure. This constrained structure is further rigidified by an extensive network of internal hydrogen bonds. Thus, the structure of SFTI-1 in solution resembles the protease-bound structure, reducing the entropic penalty upon protease binding. When cleaved at the scissile bond, it is thought that the rigidifying features of SFTI-1 maintain its structure, allowing the scissile bond to be reformed. The lack of structural plasticity for SFTI-1 is proposed to favour initial protease binding and continued occupancy in the protease active site, resulting in an equilibrium between cleaved and uncleaved inhibitor in the presence of protease. We have determined, at 1.15 Å resolution, the x-ray crystal structures of complexes between human kallikrein-related peptidase 4 (KLK4) and SFTI-FCQR(Asn14), and between KLK4 and an acyclic form of the same inhibitor, SFTI-FCQR(Asn14)*[1,14]*, with the latter displaying a cleaved scissile bond. Structural analysis and MD simulations together reveal the roles of altered contact sequence, intramolecular hydrogen bonding network and backbone cyclization, in altering the state of SFTI’s scissile bond ligation at the protease active site. Taken together, the data presented reveal insights into the role of dynamics in the standard-mechanism inhibition, and suggest that modifications on the noncontact strand may be a useful, underexplored approach for generating further potent or selective SFTI-based inhibitors against members of the serine protease family.

## Introduction

Proteases play fundamental roles in a wide range of physiological processes, including blood coagulation, complement activation, apoptosis and inflammation. Proteases are tightly regulated and excessive proteolysis is commonly associated with a variety of pathological conditions [1]. Therefore correct temporal and spatial protease activity is crucial and is regulated by numerous naturally occurring protease inhibitors and allosteric control mechanisms.

Sunflower trypsin inhibitor-1 (SFTI-1), a natural product of sunflower seeds [2], is a cyclic peptide of molecular mass 1.5 kDa. Its three-dimensional structure has been determined in solution [3] and in crystal complexes with bovine β-trypsin [2], matriptase [4] and kallikrein-4 (KLK4) [5]. With only 14 amino acids, SFTI-1 is one of the smallest and most potent naturally occurring standard-mechanism (otherwise known as Laskowski-mechanism) protease inhibitors.

SFTI-1 is stabilized by an extensive internal hydrogen bond network, by backbone cyclisation, and a disulfide bridge which confers bicyclization. This rigid arrangement minimizes entropic losses upon binding and ensures more efficient binding interactions, compared to flexible ligands [6]. Upon cleavage of the scissile bond by its target protease, trypsin, SFTI-1 maintains its structure and has the bond reformed with an equilibrium of 1:9 in favour of the intact bond, thereby avoiding the fate of ordinary substrates [2]. The unique structural features of SFTI-1 make it an attractive template for the design and chemical synthesis of novel protease inhibitors with altered specificities and high potency [7–9].

The role of the SFTI-1 recognition sequence [9], backbone cyclization [3, 10], disulfide bond [10, 11] and internal hydrogen bonding network [7, 8] have been studied by monitoring their effects on inhibitory activity. For example, tuning the SFTI protease recognition sequence increased the kon of an interaction with selected proteases, but was often accompanied by unwanted increases in koff [7]. It was reasoned that by changing the SFTI recognition sequence, the internal hydrogen bonding network was weakened, leading to increased flexibility of the SFTI-derivative which increased koff. The effects of backbone cyclization have also been studied, with a 1,14-acyclic version of SFTI-1 *(SFTI-1 [1,14])* resulting in a 20-fold reduction in inhibitory activity [3]. Since SFTI-1 is a powerful scaffold from which new selectivities can be generated, a structural appreciation of the cooperative role of backbone cyclization and changed contact sequence will inform the next generation of SFTI-based inhibitors.

SFTI has been used as a platform for the development of potent and highly selective KLK4 inhibitors. There is considerable interest in developing KLK4 inhibitors because of its potential role in a variety of diseases including prostate cancer and ovarian cancer. Over-expression of KLK4 has been documented in malignant prostate, ovarian and breast tumors [12–14] and is thought to regulate cell growth both directly through PAR2 activation [15] and indirectly through regulation of proteolytic cascades [14]. Molecular modelling and sparse matrix substrate screening resulted in creation of an engineered SFTI inhibitor with enhanced potency and selectivity towards KLK4 called SFTI-FCQR [9]. A second generation derivative, with a proposed improved intramolecular hydrogen bond network conferred by an Asp→Asn substitution at position 14, SFTI-FCQR(Asn14), resulted in a 125-fold increased inhibitory activity towards KLK4 [8]. These studies show that selective inhibitors can be generated by changing contact residues and concomitantly, potency can be increased by changing noncontact residues to restore an internal hydrogen bonding network.

In the current study we have determined crystal structures of KLK4 in complex with SFTI-FCQR(Asn14) and an acyclic form of the same inhibitor, SFTI-FCQR(Asn14)[1,14]. These structures provide an insight into the roles of altered contact sequence, the internal hydrogen bond network and backbone cyclisation in KLK4 inhibition.

## Materials and Methods

### Protein expression, refolding and purification

For crystallographic studies and the proteolysis LCMS assay, KLK4 was expressed in *E. coli* using the previously reported pET12-proPSA-hK4 chimera plasmid [16], refolded from inclusion bodies, and purified as previously described [5]. For the inhibition assay, recombinant pro-KLK4 was expressed in stably transfected *Spodoptera frugiperda* Sf9 cells and purified by nickel agarose affinity chromatography, as previously described [15]. The KLK4 pro-sequence was cleaved using thermolysin to generate mature KLK4, followed by anion exchange chromatography to isolate KLK4 from the activation reaction, as recently reported [7].

### Synthesis of SFTI variants

For crystallographic and LCMS studies, SFTI-FCQR(Asn14)[1,14] was obtained from GLS Peptide Synthesis (Shanghai, China). For all other studies, SFTI-FCQR(Asn14)[1,14] was produced using microwave-assisted solid phase peptide synthesis on 2-chlorotrityl resin. The linear peptide chain was assembled using a CEM Discover SPS Microwave system and Fmoc chemistry (full details provided in [7]), followed by cleavage from the resin and removal of side chain protection groups using 95% TFA, 2.5% thioanisole, 1.25% triisopropylsilane, 1.25% H2O. The acyclic, reduced peptide was purified by reverse-phase HPLC, then the intramolecular disulfide bond was formed by incubation in 0.15 M Tris-HCl pH 8.0, 1 mM EDTA, 10 mM reduced glutathione, 1 mM oxidized glutathione for 48 hours. The acyclic, disulfide-bridged peptide was subsequently isolated following a second round of reverse-phase HPLC, and validated by MALDI-TOF/MS.

### Inhibition assay

Inhibition of KLK4 by acyclic SFTI-FCQR(Asn14) was assessed by determining the inhibition constant (Ki) in competitive inhibition assays. A serial dilution of inhibitor was incubated with KLK4 in 96-well low-binding assay plates (Corning) and allowed to reach equilibrium. Dilutions of inhibitor and KLK4 were prepared in assay buffer (0.1 M Tris-HCl pH 7.4, 0.1 M NaCl, 0.005% Triton X-100). A constant amount of substrate (FVQR-pNA) was subsequently added to each well, and substrate cleavage was monitored by measuring the change in absorbance at 405 nm using a microplate spectrophotometer (10 s reading interval for 7 min). The final concentrations of KLK4 and FVQR-pNA were 1.5 nM and 120 μM, respectively. The *Ki* value was determined by non-linear regression (Morrison method) in GraphPad Prism 7 using activity data from three assays performed in triplicate.

### Crystallization

All crystals were grown using the hanging drop vapor diffusion method, with 1:1 (v/v) ratio of protein to mother liquor. 20 mg/mL KLK4 was incubated overnight with 3 fold molar excess of inhibitor (solubilized in 25% acetonitrile to the working concentration of 100 mM) at 4°C. Crystals were obtained within a week by mixing 2 μL of protein-peptide solution with equal volume of crystallization buffer (0.1 M Li2SO4, 0.1 M sodium acetate, 30 % PEG 8000, pH 4.6) and equilibrated at 18°C.

### X-ray data collection, structure determination and refinement

All datasets were collected at the Australian Synchrotron, Victoria, Australia on the MX2 beamline equipped with an ADSC Quantum 315r Detector [17]. Structure determination and refinement were performed as previously described [5]. Molecular replacement was performed using Phaser [18] within the CCP4 suite [19]. The modified KLK4-PABA structure with PDB ID 2BDG [20] was used as a search model. The electron densities were visualized in Coot [21] and revealed the presence of inhibitor molecules bound to the active site of KLK4. Inhibitors were manually built into the positive electron density. The models were then improved in several cycles of refinement using the program phenix.refine [22]. Model building between the cycles of refinement was carried out in Coot [21]. Structures were refined using anisotropic displacement parameters (ADP) and the default weights for stereochemical and ADP restraints were optimized automatically. Ordered solvent was added automatically and water molecules were then checked manually in Coot before the final round of refinement.

### Computational resources

Calculations, modeling and simulations were performed on MonARCH (NVIDIA GPU cluster, Monash University).

### Atomic coordinates, modeling and graphics

In MD simulations, atomic coordinates were obtained from the following PDB entries: 4K1E and 4KEL (SFTI-FCQR(Asp14) and SFTI-FCQR(Asn14) respectively). Missing residues and atoms were rebuilt using MODELLER version 9.10 [23]. Acyclic SFTI-peptide variants were modelled from the above PDBs using PyMOL version 1.8.2 [24] to introduce new N- and C-termini. All structural representations were produced using PyMOL version 2.2.1 [24] and VMD 1.9.3 [25], and all trajectory manipulation and analysis was performed with a combination of custom scripts, MDTraj [26], SciPy [27], Matplotlib [28], iPython [29] and VMD 1.9.3 [25].

### Molecular dynamics (MD) systems setup and simulation

For each protein, residue protonation states were set as appropriate at pH 7.0 using PROPKA [30, 31]. Each protein was parameterized using the AMBER ff14SB all-atom force field [32–34], placed in a rectangular box with a border of at least 12 Å of explicit TIP3P water [35] on all sides of the protein, and the system charge was neutralized by addition of sodium or chloride counter-ions.

Systems were relaxed with 15000 steps of energy minimization, followed by equilibration. In equilibration, atoms’ initial velocities were randomly distributed according to a Maxwell – Boltzmann distribution at 100 K. Harmonic positional restraints of 100 kcal^−1^ mol^−1^ Å^−2^ were applied to protein backbone atoms and temperature was steadily increased from 100 K to 300 K over the course of 100 ps, with a Langevin damping coefficient of 5 ps^−1^. Pressure was then equilibrated to 1 atm with a Berendsen barostat [36] (τ_p_ = 0.1 ps) and restraints were removed steadily over 200 ps.

Production simulations were performed in the NPT ensemble without positional restraints, using an integration timestep of 2 fs, and saving snapshots every 10 ps for analysis. Three independent replicates of each system were simulated for 500 ns each. All simulations were performed using AMBER16 [37] with periodic boundary conditions, long-range interactions were computed using PME [38].

## Results and Discussion

To discuss monocyclic and acyclic permutants of SFTI peptides, we herein adopt the nomenclature SFTI-1[A,B][C,D], where ABCD indicate the free [N,C] peptide termini in the monocyclic/acyclic peptide [39]. [6,5] indicates a peptide cleaved at the scissile bond, while [1,14] indicates a linearised peptide.

### Additional inhibitory potency gained from Asp14Asn mutation is lost upon 1,14-acyclisation

SFTI-1 is a commonly used scaffold for the development of selective protease inhibitors. Importantly, SFTI-1 functions as a standard-mechanism protease inhibitor: its rigid structure stabilises the conformation of the binding residues, preventing structural changes that would normally accompany proteolysis [2].

To compare the effect of cyclisation in the SFTI-FCQR(Asn14) peptide to previous studies on the role of cyclisation in the SFTI-1 scaffold, we determined the proteolytic inhibition constants for the bicyclic SFTI-FCQR(Asn14) inhibitor, and monocyclic SFTI-FCQR(Asn14)[1,14] inhibitor for KLK4. Monocyclic SFTI-FCQR(Asn14)[1,14] was found to inhibit KLK4 activity with a Ki of 3.48 nM compared to 0.04 nM for the bicyclic SFTI-FCQR(Asn14) (**Figure 1**).

**Figure 1.**
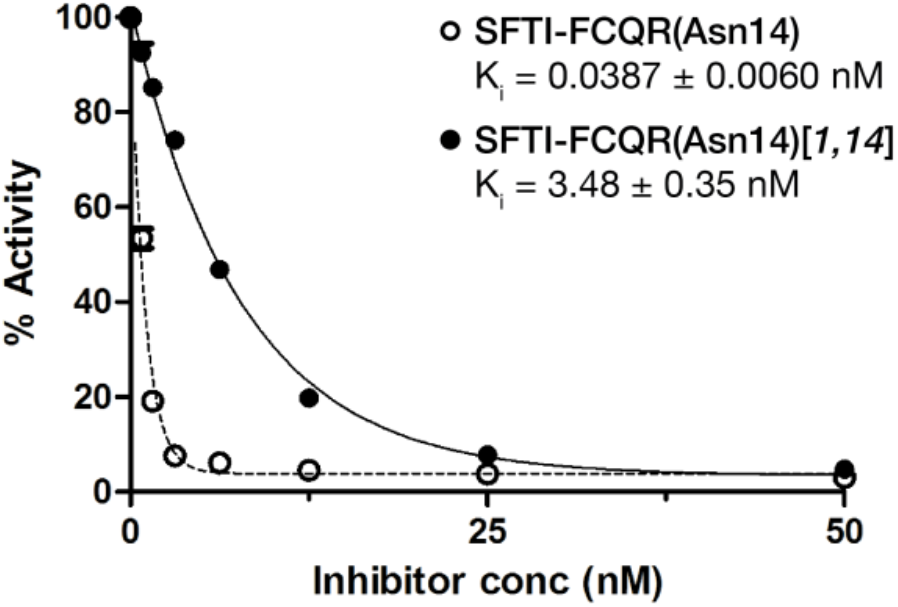
Inhibition of KLK4 by SFTI-FCQR(Asn14) cyclic and acyclic permutants.

The rigidity of SFTI-1 is believed to extend to its cleaved form, SFTI-1*[6,5]*, maintaining its structure and ability to bind trypsin. Trypsin is able to religate the scissile bond of SFTI-1[6,5], resulting in a 9:1 equilbrium between SFTI-1:SFTI-1[6,5] after 2 hours, favouring the bicyclic variant [39]. Backbone cyclisation is a prominent feature of SFTI-1 that assists in its potency as a protease inhibitor, though it is not essential. As such *1,14-*acyclisation does not prevent SFTI-1 from achieving scissile bond resynthesis. SFTI-1[1,14], when incubated with trypsin, shows very slow cleavage to SFTI-1*[1,14][6,5]*, also resulting in an equilibrium between SFTI-1*[1,14]* and SFTI-1*[1,14][6,5]* after 2 hours [10]. More recently, a study incubated trypsin with SFTI-*1[1,14][6,5]* and by a 3 hour timepoint, the main product was a religated SFTI-1*[6,5]* [40].

It is reasonable to expect that all derivatives of SFTI-1 are able to achieve some level of scissile bond religation. In line with previous studies on SFTI-1, *1,14*-acyclisation is not expected to dramatically affect the structure of the SFTI canonical loop [3], though weaker inhibition compared to bicyclic SFTI variants is expected as a result of an increased entropic debt on binding [3, 10].

The Asp14Asn mutation was designed on an entropic rationale: to strengthen the internal H-bond network within SFTI-FCQR, and further rigidify the bicycle [8]. It is interesting to note that the SFTI-FCQR(Asn14)[1,14] variant is as potent as the bicyclic SFTI-FCQR(Asp14) [9] (Ki=3.48 nM, and 3.59 nM, respectively). This suggests that the Asp14Asn mutation increases inhibitory potency as much as the 1,14-acyclised backbone reduces it, and by extension, the Asp14Asn mutation should decrease the entropic debt of binding as much as acyclisation increases it.

### Structural rationalisation for potency of SFTI-FCQR(Asn14) against KLK4

In a previous study, we determined the crystal structure of SFTI-FCQR(Asp14) in complex with KLK4, and found that improvement of KLK4 inhibition by SFTI-FCQR(Asp14) over the wild type SFTI-1 is primarily due to the optimised contact residues, which improve complementarity between P4 Phe and the hydrophobic S4 subsite, P2 Gln and the S2 subsite, and increased interactions between P1 Arg and the S1 subsite [5].

In order to investigate the increased potency of the related peptide SFTI-FCQR(Asn14) over SFTI-FCQR(Asp14), we determined the crystal structure of SFTI-FCQR(Asn14) in complex with KLK4 at 1.15 Å resolution (**Table 1**). The asymmetric unit consists of one KLK4 molecule bound to one molecule of SFTI-FCQR(Asn14) inhibitor (**Figure 1b**). The mean temperature factor for the inhibitor atoms is 14.9 Å^2^, compared with the average of 10.8 Å^2^ for the atoms of KLK4. The electron density around the scissile bond (**Figure 1a**) is well defined for the SFTI-FCQR(Asn14) inhibitor, confirming that it is present within the crystal in an uncleaved state bound to the protease (**Figures 1b, 1c**). Electron density for loop III backbone atoms was sufficient for modelling, but density corresponding to loop III side chains E74, D75, Q76, E77 was not detected. This observation is identical to the previously reported structure of KLK4-SFTI-FCQR(Asp14) [5]. It is worth noting that this loop is involved in a crystal contact with residues 10-12 of chain B (the SFTI-derivative) in a symmetry partner (−X, Y+1/2, −Z) + (−1, 0, 0).

**Table 1.** Data collection and refinement statistics for the KLK4-FCQR(Asn14) and KLK4-FCQR(Asn14)[2,24][6,5] structures.

|  | KLK4–FCQR(Asn14) | KLK4–<br>FCQR(Asn14)[1,14][6,5] |
| --- | --- | --- |
| **Data collection** |  |  |
| Source | Australian Synchrotron MX2<br>Beamline | Australian Synchrotron MX2<br>Beamline |
| Wavelength (Å) | 0.9537 | 0.9537 |
| Space group | P12<sub>1</sub>1 | P12<sub>1</sub>1 |
| Unit cell dimensions (Å) | a=39.84, b=63.33, c=41.19 | a=39.53, b=63.15, c=40.89 |
| Angles (°) | $\alpha=\gamma=90$, $\beta=115.58$ | $\alpha=\gamma=90$, $\beta=114.92$ |
| Resolution (Å) | 37.15–1.15 | 37.09–1.15 |
| Number of measured reflections | 268420 | 247290 |
| Number of unique reflections | 64890 | 62317 |
| Completeness (%) | 99.0 (98.1) | 96.6 (93.6) |
| Redundancy | 4.1 (3.5) | 4.0 (3.5) |
| R<sub>pim</sub> (%) | 3.9 (22.9) | 4.0 (27.8) |
| CC1/2 | 0.997 (0.855) | 0.998 (0.883) |
| <I/σ(I)> | 10.8 (4.3) | 13.0 (3.0) |
| **Molecular replacement** |  |  |
| Number of molecules in the asymmetric unit | 1 | 1 |
| **Refinement** |  |  |
| Number of protein atoms | 1875 | 1870 |
| Number of water molecules | 209 | 238 |
| R<sub>work</sub> (%) | 13.66 | 12.98 |
| R<sub>free</sub> (%) | 16.34 | 16.35 |
| *Rms deviations from ideality* |  |  |
| Bond lengths (Å) | 0.006 | 0.007 |
| Bond angles (°) | 1.218 | 1.044 |
| Average B factors (Å<sup>2</sup>) |  |  |
| Protein | 11.3 | 11.02 |
| Solvent | 22.92 | 23.51 |
| *Ramachandran plot* |  |  |
| Favoured (%) | 98.00 | 98.65 |
| Allowed (%) | 100.0 | 100.0 |
| Outliers (%) | 0 | 0 |
| MolProbity score | 0.52, 100<sup>th</sup> percentile | 1.09, 98<sup>th</sup> percentile |
| PDB ID | 4KEL | 6O21 |

<sup>a</sup>Values in parentheses refer to the highest resolution shell.
<sup>b</sup>Agreement between intensities of repeated measurements of the same reflections and can be defined as: $\sum(I_{h,i} - \langle I_h \rangle) / \sum I_{h,i}$ , where $I_{h,i}$ are individual values and $\langle I_h \rangle$ is the mean value of the intensity of reflection $h$ .

The interaction between protease and inhibitor is dominated by the P1 residue Arg5 which makes six hydrogen bonds and three salt bridges with KLK4. Six additional hydrogen bonds are formed between SFTI-FCQR(Asn14) backbone atoms Gly1, Cys3, Gln4, Ser6 and Ile7 and KLK4 (**Figure 1d, Supp Table 1**). These interactions are identical to those in the KLK4-SFTI-FCQR(Asp14) complex [5].

The internal H-bond network of SFTI is hypothesized to make an important contribution to its binding kinetics (increased k_on_) and prolonged occupancy (decreased k_off_). It has been suggested that this internal hydrogen bonding network can be weakened when optimizing the recognition sequences for targeting specific proteases [7, 8]. Our previously reported structure of KLK4 in complex with SFTI-1 showed 10 internal SFTI hydrogen bonds; optimising the protease contact residues from RCTK to FCQR resulted in only 6 internal hydrogen bonds remaining [5]. Mutating Asp14 to Asn14 was previously hypothesized to strengthen the hydrogen bonding network of SFTI-FCQR, contributing to the higher potency of the Asn14 variant [8]. However, the 6 internal H-bonds in KLK4-SFTI-FCQR(Asn14) (**Figure 1b**) are identical to the 6 internal hydrogen bonds in the KLK4-SFTI-FCQR(Asp14) complex [5]. Specifically, the hydrogen bond between the side chain oxygen of Asp/Asn14 and the backbone nitrogen of Phe2 is preserved (**Figure 1b**).

Thus the crystallographic evidence presented here does not provide an explanation for the increased potency of the Asn14 inhibitor (Ki of 0.04 nM [8] vs 3.59 nM [9]).

### Serendipitous observation of a cleaved peptide bound to the protease active site

To determine the structural effects of backbone acyclisation on this SFTI derivative when bound to a protease, we solved the crystal structure of a monocyclic SFTI-FCQR(Asn14)[1,14] inhibitor in complex with KLK4. The crystals are isomorphous with those of the complex of KLK4 with the bicyclic variant, and diffracted to a resolution of 1.15 Å (**Table 1**). The overall structure of the protease is largely unchanged, compared to KLK4-SFTI-FCQR(Asn14), with an RMSD of 0.54 Å over all non-hydrogen atoms (Figure 4a).

In contrast, the structure of the inhibitor is dramatically different from the bicyclic variant (Figure 3a). Electron density at 1σ of a well-defined neo-C terminus after the inhibitor’s Arg5 shows this peptide bond has been cleaved, producing a fully acyclic structure with only the disulfide bond connecting the fragments. There is also a lack of electron density for the residues of the new amino terminus (Ser6-Pro8) and the side chain of Asn14 such that these residues / atoms could not be modelled (Figures 3b, 3c). Thus incubating KLK4 with monocyclic SFTI-FCQR(Asn14)*[1,14]* over the course of crystallisation (5 days) has serendipitously trapped a substrate cleaved at the scissile bond (Arg5ļSer6). Acyclisation and further cleavage is consistent with greater observed flexibility—the 1,14-acyclic 6,5-cleaved structure is more conformationally dynamic with a mean temperature factor for the inhibitor atoms of 20.1 Å^2^, compared to a temperature factor of 14.9 Å^2^ for the bicyclic version and compared to an average of 11.3 Å^2^ for the atoms of KLK4 within the KLK4-SFTI-FCQR(Asn14)*[1,14][6,5]* complex.

**Figure 2.**
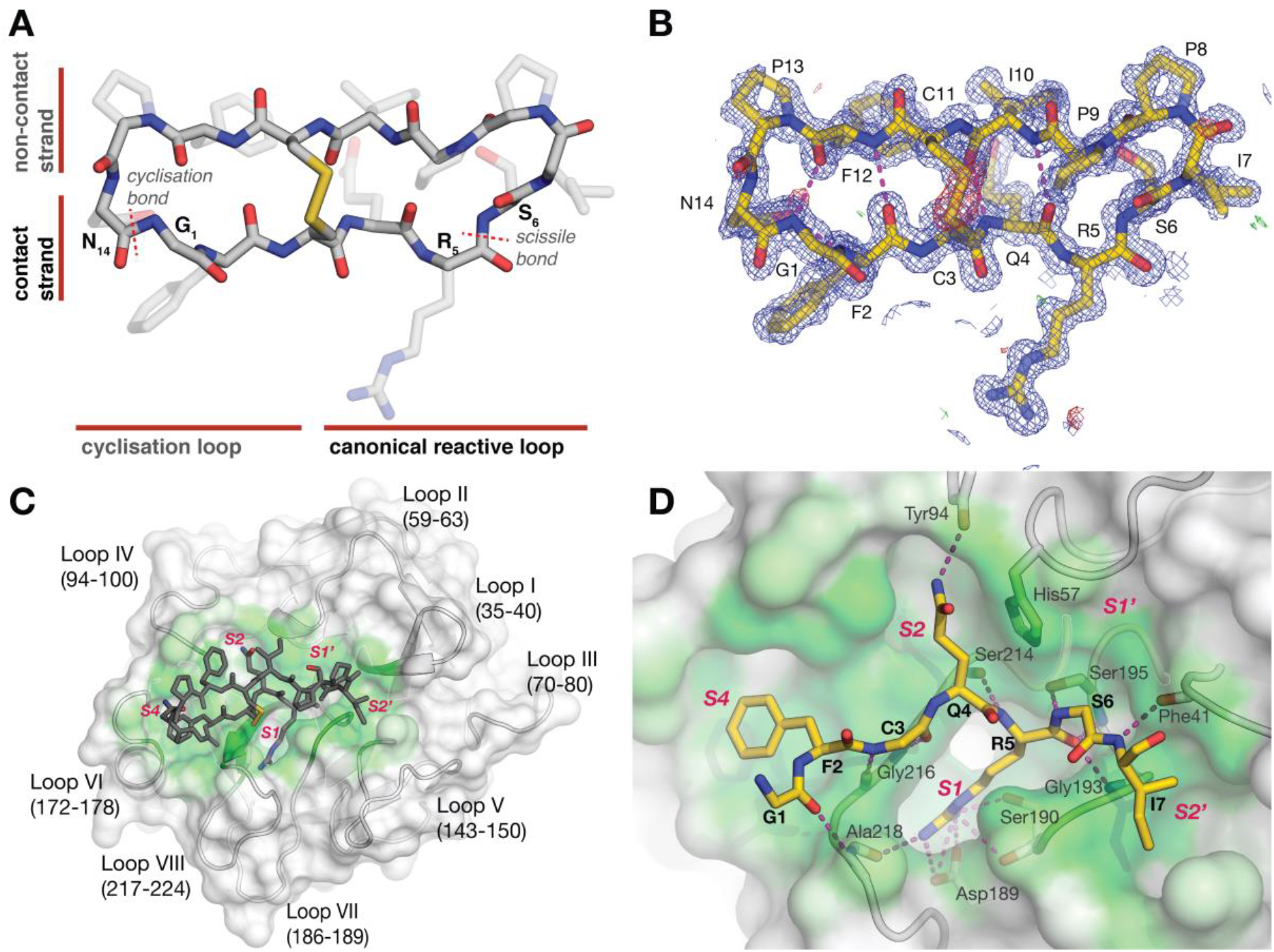
**(A)** Structural features of a bicyclic SFTI derivative. SFTI-FCQR(Asn14) is shown as grey sticks, with a solid backbone and transparent sidechains. The upper β-strand is the “noncontact strand”, and the lower is the “contact strand”, containing the specificity determining residue. The left “cyclisation loop” contains a peptide bond between residues 14 and 1. The “reactive loop” on the right contains the scissile bond between residues 5 and 6. **(B)** Electron density within SFTI-FCQR(Asn14). The inhibitor SFTI-FCQR(Asn14) from the crystal structure in complex with KLK4 has been taken out of the context of KLK4 to show the intramolecular hydrogen bond network. SFTI-FCQR(Asn14) is shown as yellow sticks, with a solid backbone, and translucent sidechains. Intramolecular hydrogen bonds are shown as magenta broken lines. Within 2.0 Å of the displayed atoms, 2*m*Fo-*D*Fc electron density contoured at 1σ, is shown as a blue mesh, *m*Fo-*D*Fc at 3σ is shown as red and green mesh. **(C)** Overall interaction between KLK4 and SFTI-FCQR(Asn14). KLK4 is shown as a cartoon, with a surface depicting surface burial by SFTI: from white (completely solvent-exposed) to green (completely buried surface). SFTI-FCQR(Asn14) is shown as grey sticks. **(D)** Intermolecular interactions between KLK4 and SFTI-FCQR(Asn14) contact strand. KLK4 is shown as a cartoon & buried surface as previously described. SFTI-FCQR(Asn14) contact strand (residues 1-7) is shown as gold sticks, with sidechains of Cys3 and Ser6 hidden for clarity. Hydrogen bonds between KLK4 and SFTI are shown as magenta broken lines.

**Figure 3.**
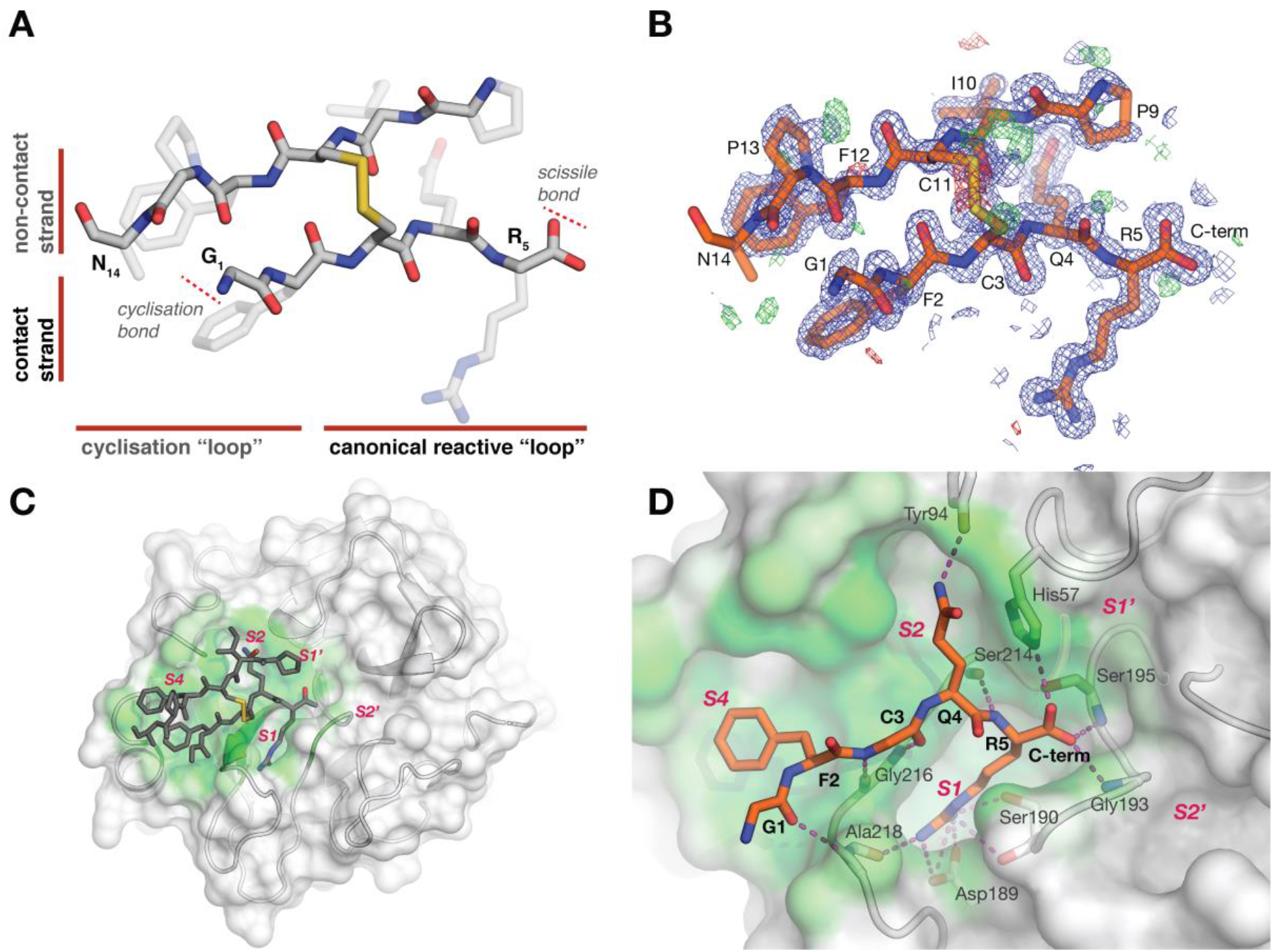
**(A)** Structural features of a cleaved, acyclised SFTI derivative. SFTI-FCQR(Asn14) [1,14] [6,5] is shown as grey sticks, with a solid backbone and transparent sidechains. The upper β-strand is the “non-contact strand”, and the lower is the “contact strand”, containing the specificity determining residue. The left “cyclisation loop” does not have a peptide bond between residues 14 and 1. The “reactive loop” on the right has been cleaved between residues 5 and 6, and displays a carboxylate group at the C-terminus of Arg5. (B) Electron density within SFTI-FCQR(Asn14)*[1,14][6,5]*. The inhibitor SFTI-FCQR(Asn14)*[1,14][6,5]* from the crystal structure in complex with KLK4 has been taken out of the context of KLK4 to show the intramolecular hydrogen bond network. SFTI-FCQR(Asn14)*[1,14][6,5]* is shown as orange sticks, with a solid backbone, and translucent sidechains. Intramolecular hydrogen bonds are shown as magenta broken lines. Within 2.0 Å of the displayed atoms, 2*m*Fo-*D*Fc electron density contoured at 1σ, is shown as a blue mesh, *m*Fo-*D*Fc at 3σ is shown as red and green mesh. **(C)** Overall interaction between KLK4 and SFTI-FCQR(Asn14)*[1,14][6,5]*. KLK4 is shown as a cartoon, with a surface depicting surface burial by SFTI: from white (completely solvent-exposed) to green (completely buried surface). SFTI-FCQR(Asn14)*[1,14][6,5]* is shown as grey sticks. Residues Ser6, Ile7 and Pro8 and part of Asn14 are not visible in the electron density of the complex. **(D)** Intermolecular interactions between KLK4 and SFTI-FCQR(Asn14)*[1,14][6,5]* contact strand. KLK4 is shown as a cartoon & buried surface as previously described. SFTI-FCQR(Asn14)*[1,14][6,5]* contact strand (residues 1-7) is shown as orange sticks, with the sidechain of Cys3 hidden for clarity. Hydrogen bonds between KLK4 and SFTI are shown as magenta broken lines.

### Interactions between SFTI-FCQR(Asn14)[1,14][6,5] and KLK4

While the monocyclic SFTI-FCQR(Asn14)*[1,14]* inhibitor has 87-fold reduction of inhibitory activity compared to bicyclic SFTI-FCQR(Asn14), it still inhibits KLK4 with a Ki of 3.48 nM. To rationalise this on a structural basis, we examined the hydrogen bonding and salt bridge formation for the complex beween KLK4 and SFTI-FCQR(Asn14)*[1,14][6,5]*. A retention of many of the interactions seen in the bicyclic structure explains how the P1-P1’ cleaved inhibitor can still be retained at the KLK4 active site. 11 hydrogen bonds and 3 salt bridges are observed between this 1,14-acyclic 6,5-cleaved peptide and KLK4 compared to the 12 hydrogen bonds and 3 salt bridges for the bicyclic structure (**Figure 3d**). Hydrogen bonds contributed by Gly1, Cys3, Gln4 and Arg5 are all preserved, as are the three salt bridges from Arg5 (see **Supp Table 1**). Thus while significant differences in the bound surface between the 1,14-acyclic 6,5-cleaved structure are seen compared to the bicyclic structure, the presence of these interactions is consistent with its inhibitory potency as well as its retention in the active site after P1-P1’ hydrolysis.

The 87-fold reduction in Ki compared to bicyclic SFTI-FCQR(Asn14) is mirrored by a reduction in the interactions with KLK4. In the SFTI-FCQR(Asn14)*[1,14][6,5]* structure, P1’-P3’ residues Ser6, Ile7 and Pro8 have no observed electron density, which we interpret as evidence that they are not engaged in stable interactions with KLK4 as they were in the bicyclic structure. Additionally, the lack of density for the side chain of Asn14 (**Figure 3b**) shows that this region remains flexible when bound to KLK4, as observed in previous studies [2, 5]. It is apparent that the Asn14 sidechain is not participating in stable hydrogen bonding within this crystal structure that would either restrict inhibitor ejection or stabilise the overall cleaved inhibitor structure.

### The role of “non-contact strand” residues in binding after scissile-bond cleavage

An alignment of the protease heavy atoms between the presented structures shows relatively large deviations of the binding pose between the SFTI-FCQR(Asn14)*[1,14][6,5]* inhibitor and the bicyclic SFTI-FCQR(Asn14) variant (**Figure 4a**). The inhibitor molecules show the smallest deviations for the contact residues 2FCQR5 (sub-1 Å differences), consistent with the retention of many of the hydrogen bonds and salt bridges between KLK4 and P1-P4. In contrast, significant differences are seen in the non-contact strand, which shows backbone displacements ranging from 2.2 Å at Pro13 to 4.8 Å at Pro9. Even greater shifts are observed for the sidechains of the non-contact sheet / strand. The sidechain of Phe12 is shifted by 10 Å and greater than 4 Å shifts are seen at Pro9 and Ile10. Since Pro9 is the first prime side residue seen in the cleaved structure (P4’) and is displaced by almost 5 Å, this suggests the unobserved residues P1’–P3’ have been ejected from the protease and have moved far from their equivalent positions in the bicyclic structure.

**Figure 4.**
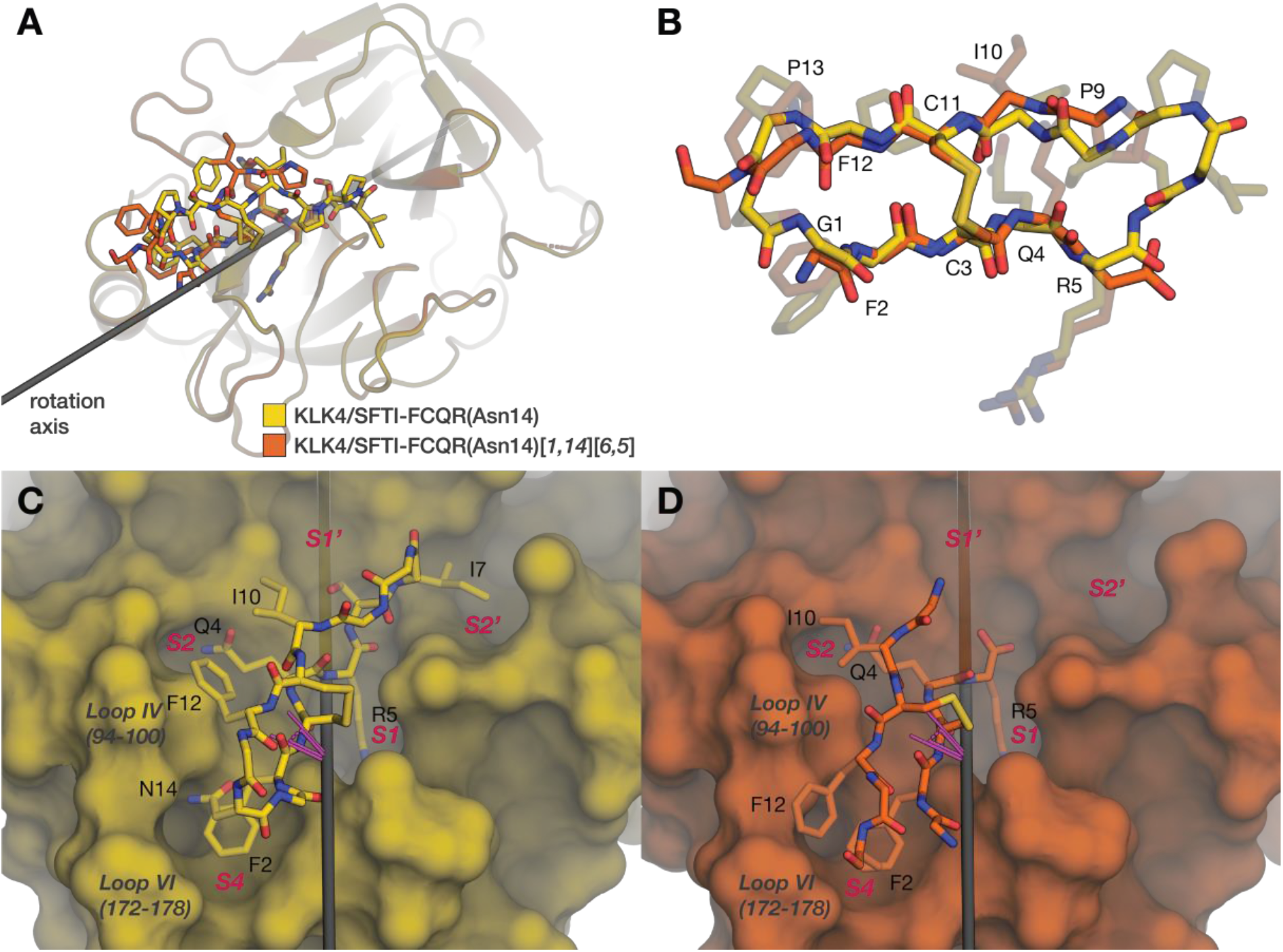
**(A)** Overall comparison between the KLK4/SFTI-FCQR(Asn14) complex (yellow), and the KLK4/SFTI-FCQR(Asn14)[1,14][6,5] (orange), after alignment on protease heavy atoms. The best-fit axis of rotation between the inhibitors is shown as a grey rod. **(B)** Comparison between the two inhibitors, after alignment on inhibitor backbone atoms. Residues which are observed in both crystal structures are labelled. **(C, D)** A side-by-side comparison of inhibitor poses within complexes. Proline side-chains are hidden for clarity; inhibitor backbone is outlined. Best-fit axis of rotation is shown as a grey rod, and a rotation angle in magenta.

Despite large changes in the inhibitor pose in the protease active site, an alignment between the backbones of the inhibitor molecules shows comparatively smaller internal deviations between the two structures (**Figure 4b**). The major differences here are in the positioning of Asn14 and the conformation of Pro9, which show maximum shifts of 6.2 Å and 2.3 Å respectively; in comparison, the backbone atoms of the remaining residues reveal a highly rigid structure between the β-turn regions. This rigidity is somewhat explained by a comparison of the internal hydrogen bonding network of the cleaved substrate within this structure (**Figures 2b, 3b**). Compared to 6 internal hydrogen bonds for the bicyclic peptide bound to KLK4, there are 4 internal hydrogen bonds within the SFTI-FCQR(Asn14)*[1,14][6,5]* peptide. Hydrogen bonds involved in the 14-1 β-turn (Gly1N-Phe12O and Phe2N-Asn14Oô) are lost in favour of Phe2N-Phe12O, parallelling previous observations in NMR solution structures of SFTI-1 [3]. Towards the scissile bond, the internal H-bond Ile10N-Gln4O is replaced by Ile10N-Gln4Oε, and the absence of Ser6-Ile7 from the acyclic structure suggests that the Ser6Oγ-Pro8O bond is lost.

Encouraged by the rigidity of the SFTI scaffold core, we examined the rigid-body motion of the SFTI backbone between the two structures, finding a 0.4 Å translation coupled with a 23.4° rotation (**Figure 4c, 4d**). This large rotation of the SFTI within the active site is driven by changes in two distinct areas between SFTI and KLK4. Firstly, contacts between Ile7 and the S2’ site are lost as a result of N-terminal ejection. Secondly, new compensatory interactions are formed with the “non-contact strand”: Phe12 and the top of loop VI, as well as Ile10 and loop IV (**Figures 4c, 4d, Supp Table 2**). A previously determined structure for trypsin-SFTI-1[1,14] (PDB:4XOJ) [40] shows that 1,14-acyclisation alone does not necessitate repositioning of SFTI-1 within the binding cleft, strongly suggesting that the rotation we observe with SFTI-FCQR(Asn14)[1,14][6,5] is a result of cleavage at the scissile bond.

### MD simulations of [6,5]-cleaved SFTI variants reveal a diverse conformational ensemble and increased flexibility in the oxyanion pocket

To examine how cleavage of the scissile bond affects the conformations accessible to SFTI, we performed molecular dynamics (MD) simulations of various KLK4-SFTI complexes. We simulated complexes with the intact SFTI-FCQR and SFTI-FCQR(Asn14) peptides, as well as these peptides cleaved at the 6,5 scissile bond, or with free termini at the 1,14 site of backbone cyclisation. Simulations were performed for 500 ns to allow the SFTI molecule sufficient time to explore alternative binding poses that may be relevant for its potency.

All simulations were stable over 500 ns, with all-atom RMSDs reaching a plateau of 3.5-4.5 Å after 25 ns (**Supp Fig 1**). In all simulations, the SFTI derivative was securely bound to the S1 pocket, with the Arg5-Asp189 salt bridge permanently occupied (**Supp Fig 4**). Simulations of complexed *[6,5]*-cleaved inhibitors showed that these were by far the most flexible variant of the SFTI peptide. As some SFTI-binding poses were only visited once across the triplicate simulations of *[6,5]*-cleaved inhibitors, we believe that these simulations represent only a small region of the complete conformational landscape of *[6,5]*-cleaved SFTI-inhibitors bound to KLK4, and that alternative methods are required to adequately sample the conformational space accessible to *[6,5]*-cleaved SFTI inhibitors. Increased conformational dynamics was mainly restricted to the inhibitor molecule (**Supp Fig 3**), though we observed a slight, but consistent increase in backbone dynamics of the oxyanion pocket loop Cys191-Pro198 in all simulations with the *[6,5]*-cleaved peptides (**Supp Fig 2**). Both SFTI variants explore the tilted binding pose observed in the KLK4-SFTI-FCQR(Asn14)[1,14]*[6,5]* crystal structure, highlighting that this is a result of *[6,5]*-cleavage, and not *[1,14]*-acyclisation. We observed that the tilted binding pose only occurs after ejection of the neo-N terminus.

In an attempt to rationalise the increased potency of the Asn14 mutation, we compared simulations of bicyclic inhibitors SFTI-FCQR and SFTI-FCQR(Asn14). The Asn14 mutation was designed in an effort to strengthen the internal hydrogen bonding network of SFTI-FCQR [8], and it appears that this was successful. We observed that Asn14 was able to make a hydrogen bond to the backbone at Phe2N in the bicyclic simulations with a mean occupancy of 21.0% (3.8%) when bound, and 40.5% (6.2%) when free, as opposed to <2% with Asp14 *(reported as mean (sd)).* Overall, the SFTI-FCQR(Asn14) inhibitor showed 3.38 (0.44) mean internal H-bonds when free, and 3.96 (0.19) mean internal H-bonds when bound to KLK4; compared to the SFTI-FCQR(Asp14) inhibitor, which showed 2.23 (0.68) when free, and 3.61 (0.04) bound to KLK4. Conformational dynamics of KLK4, as measured by Cα root mean square fluctuations (RMSF), was no different between any of the intact simulations, nor was it significantly different between the inhibitor molecules (**Supp Fig 3a, 3d**).

### Implications for the design of SFTI-derived protease inhibitors

The rigidity of the SFTI scaffold is often asserted to be responsible for the low entropic debt of SFTI-derivatives on binding, and hence their strong potency as inhibitors [41, 42]. In highlighting the rigid-body motion of the β-sheet core upon cleavage & N-terminal ejection, however, we provide direct evidence that this β-sheet hydrogen bonding network may also serve a second purpose in maintaining the conformation of the inhibitor after scissile bond cleavage.

As far as we are aware, we report the first crystal structure of an SFTI derivative that shows significant interactions between side chains of the non-contact strand and the protease. So far, these residues have been assumed to play a minimal role in the inhibitory potency of SFTI derivatives [5], and few studies have been performed analysing the effects of mutating these residues. While this region is involved in a crystal contact, the observed packing of Phe12 and Ile10 of the cleaved SFTI against the protease in the KLK4-SFTI-FCQR(Asn14)*[1,14][6,5]* structure suggests that the non-contact strand may be useful for further tuning of inhibitory potency, or selectivity, as this packing has not been observed in other isomorphous crystals of KLK4-SFTI-derivative complexes.

### Implications for the standard-mechanism

Standard-mechanism (or Laskowski-mechanism) inhibitors interact with proteases as tight-binding substrates, and behave following the reaction mechanism

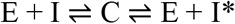

where E is the protease, I is the inhibitor, I* is the cleaved inhibitor, and C is a tight-binding complex [43].

Subsequent kinetic studies of these inhibitors have extended this mechanism to include additional stable intermediates

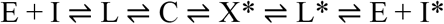

including L and L* as loose complexes, and X* as an “additional intermediate” seen on titration of the cleaved inhibitor with the protease, in which the substrate is still cleaved [44]. These studies highlight that X* is a species in which the scissile bond is completely cleaved, and not covalently bound to the protease [45, 46].

Structures of intact Laskowski (or ‘standard-mechanism’) inhibitors bound to proteases have shed light on the structure of the structure of the complex C as a Michaelis complex, with an unmodified inhibitor bound to the enzyme. As this form is almost exclusively observed in crystallised protease–inhibitor complexes, it can be confidently stated that the Michaelis complex is usually the thermodynamically favoured complex. However, crystallography and kinetics experiments have so far struggled to explain how these inhibitors resist proteolysis, and two potentially complementary mechanisms have been proposed [42, 47, 48]. Firstly, strong non-covalent bonds in the Michaelis complex have to be broken in order to proceed to the transition-state complex, making this step unfavourable. Secondly, following the formation of the acyl-enzyme intermediate, the neo-N-terminus is retained in close proximity to the acyl linkage, favouring its attack on the acyl group (and hence resynthesis of the scissile bond) over an attack by a water molecule to complete the hydrolysis [48].

The rigidity of enzyme–inhibitor complexes—and in particular, the inhibitor’s reactive loop—support both of these explanations of the slow rate of hydrolysis proceeding from the Michaelis complex [41, 42], and so it is unclear to what extent each mechanism contributes to hydrolysis resistance. In an earlier study, an initially P1-P1’-cleaved Laskowski inhibitor (bovine pancreatic trypsin inhibitor, BPTI) was crystallised with catalytically inactive trypsin-S195A [41]. In this structure, the scissile bond termini of the cleaved Laskowski inhibitor remained in close proximity to each other, as well as the (inactive) catalytic machinery, in a conformation described as ‘poised to religate’, providing support for the second mechanism.

Our crystal structure of KLK4-SFTI-FCQR(Asn14)[1,14]*[6,5]*, to the best of our knowledge, represents the first cleaved standard mechanism inhibitor bound to a catalytically active protease. Given that it was formed on incubating a scissile-bond intact inhibitor with the protease, it appears that in spite of the rigidifying effects of the Asn14 mutation [8], the [1,14]-acyclisation and Thr4Gln substitution have weakened the internal hydrogen-bonding network, perhaps beyond a critical threshold, to shift the thermodynamically favoured intermediate beyond the Michaelis complex C, to a cleaved form X*. It is important to note that SFTI-FCQR(Asn14)[1,14] is still a potent inhibitor with a Ki of 3.48 nM.

This structure also provides the first insight into the structure of the catalytic intermediate X*—a completely cleaved species with N-terminal ejection. Importantly, our crystal structure demonstrates correct catalytic geometry, yet it is clear that the reactive loop in monocyclic SFTI-FCQR(Asn14)[1,14] is flexible and thus unable to maintain the P1’ N-terminus in close proximity to the catalytic machinery. We therefore suggest that proteolysis resistance of SFTI-derived inhibitors does not rely alone on maintaining the position of the scissile peptide termini in close proximity. It appears that the difficulty of moving from the Michaelis complex to the tetrahedral intermediate is a significant rate-limiting factor in proteolysis of SFTI derivatives. We cautiously suggest that this might also be true for the protolysis resistance of standard-mechanism inhibitors.

## Conclusions

By determining the structures for bicyclic and monocyclic SFTI derivatives, we show how a Laskowski-mechanism inhibitor, such as SFTI, can be driven further than usual down the substrate pathway through changes in dynamics. The normal pathway for such an inhibitor starts with long-range interactions with a protease followed by direct binding in a catalytically competent conformation. Upon binding, cleavage of the P1-P1’ bond can occur, but the internal network of hydrogen and disulfide bonds in Laskowski-mechanism inhibitors provides rigidity to the inhibitor’s reactive loop. It is often asserted that this rigidity functions to maintain the N- and C-termini in close proximity, promoting efficient scissile bond religation [41, 47]. Here in SFTI-FCQR(Asn14), P4-P1 optimisation, followed by [1,14]-acyclisation, has weakened the internal hydrogen bonding network beyond a critical threshold. The reactive loop no longer maintains a rigid conformation upon cleavage, enabling prime-side ejection.

To the best of our knowledge, this is the first crystal structure of a catalytically active serine protease bound to a P1-P1’ cleaved Laskowski-mechanism inhibitor. Previous studies have only been able to show *pre-cleaved* substrates [49] or *pre-cleaved* inhibitors [41] subsequently incubated with catalytically inactive S195A variants of serine proteases. These structures allow us to dissect the forces involved in the latter stages of the protease mechanism: P1-P1’ cleavage, and product ejection. The rigidity of the reactive loop no doubt slows down the rate of successful cleavage events, and plays a significant role in the potency of Laskowski-mechanism inhibitors. Our presented structures and MD simulations, however, suggest that SFTI-based inhibitors may also be able to rely on additional interactions using residues of the “non-binding strand” to minimise koff, prolonging their inhibition after cleavage.

In this work, we have presented multiple subtle variants of a standard/Laskowski-mechanism inhibitor, and shown that these can alter the substrate:inhibitor equilibrium. This has implications for understanding how standard-mechanism inhibitors manage to re-form the scissile bond, and suggests novel sites of interactions which may be relevant to this process. It is clear that the conformational dynamics of standard-mechanism inhibitors are still not fully understood, and modifications that tune these dynamics and conditional interactions may be useful in generating further potent or selective SFTI-based inhibitors against members of the serine protease family.

## Acknowledgements

We would like to thank Dr. Peter Goettig (Universität Salzburg) for helpful comments and criticism during preparation of this manuscript. We thank the Monash eResearch Center for computational resources and assistance.

## Accession Numbers

The coordinates have been deposited in the Protein Data Bank. KLK4-SFTI-FCQR(Asn14) (4KEL); KLK4-SFTI-FCQR(Asn14)*[1,14][6,5]* (6O21).

## Abbreviations

KLK: Kallikrein-related peptidase
RMSD: Root Mean Square Deviation
SFTI: Sunflower trypsin inhibitor
SFTI-FCQR(Asn14): bicyclic SFTI substituted with Phe2, Gln4, Arg5 and Asn14
SFTI-FCQR(Asn14)*[4,14]*: monocyclic SFTI cleaved at 1,14 position
SFTI-FCQR(Asn14)*[1,14][5,6]*: acyclic SFTI cleaved at 1,14 and 5,6 positions

