## Supporting Information for "KLK4 inhibition by cyclic and acyclic peptides: structural and dynamical insights into standard-mechanism protease inhibitors"

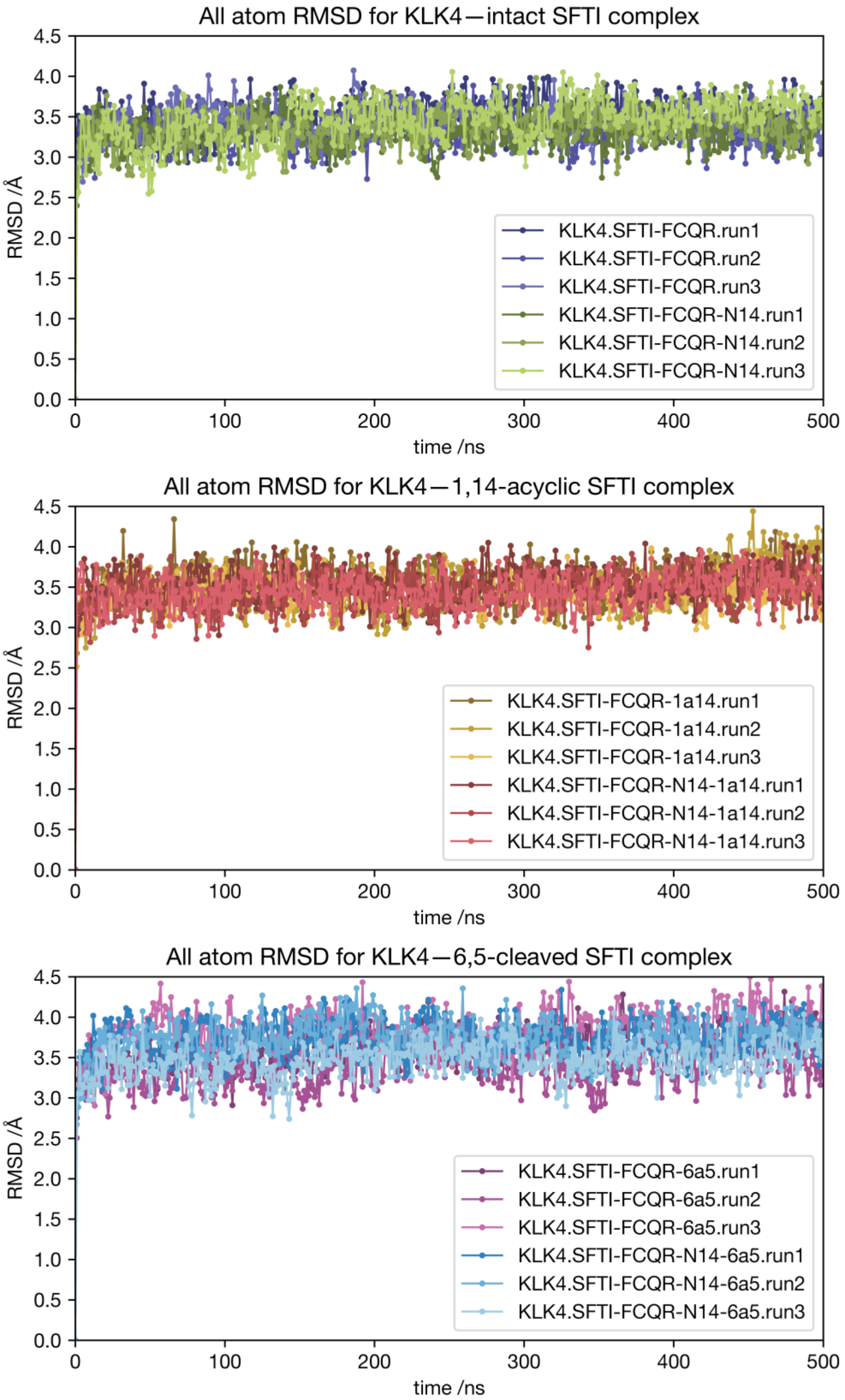


**Supporting Figure 1. All-atom RMSD of simulations.** Root mean square deviation (RMSD) plots of all protease and inhibitor atoms in **(A)** KLK4 with bicyclic SFTI simulations, **(B)** KLK4

with 1,14-acyclised SFTI-simulations, and **(C)** KLK4 with 6,5-cleaved SFTI simulations.

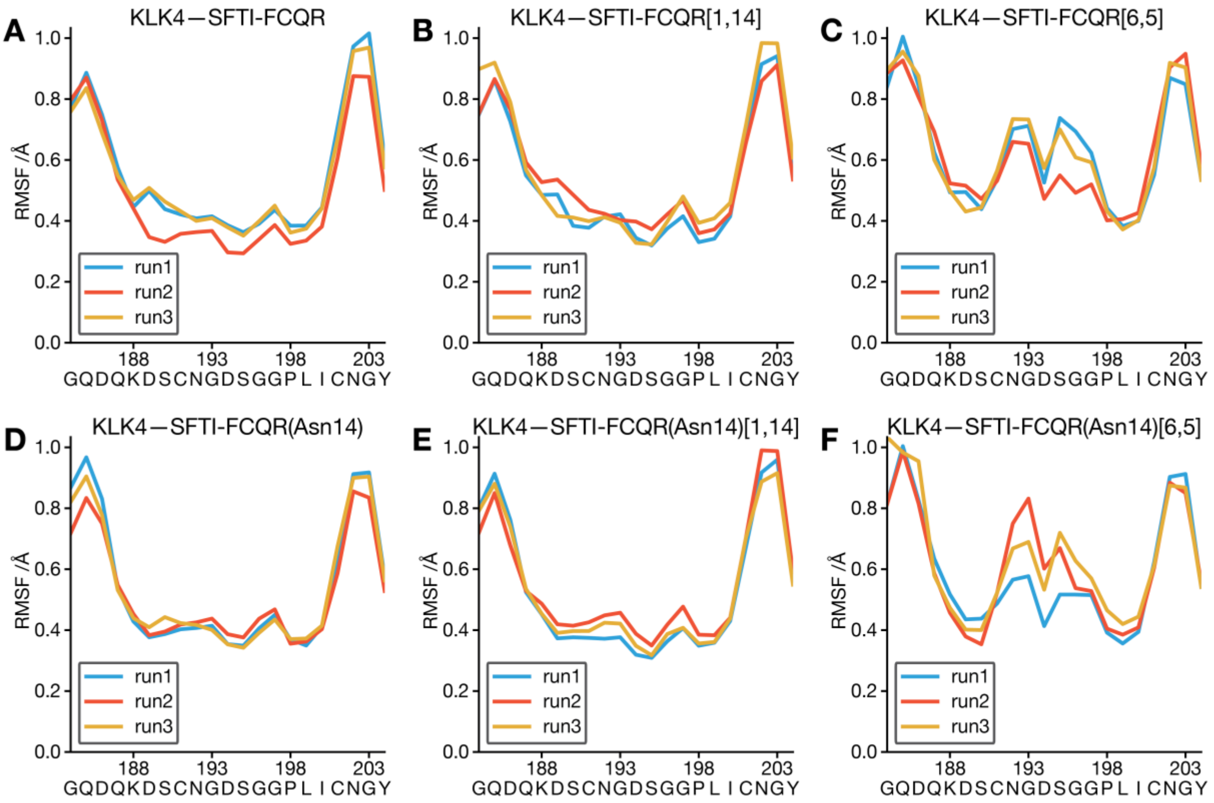


**Supporting Figure 2. Cα-RMSF of KLK4 residues 186−208.** Root mean square fluctuation (RMSF) plots of KLK4 residue 186−208 Cα atoms in **(A, B, C)** simulations of KLK4 with SFTI-FCQR, and **(D, E, F)** simulations of KLK4 with SFTI-FCQR(Asn14). Left column contains bicyclic permutants of the SFTI inhibitor, middle column contains 1,14-acyclised permutants, and right column contains 6,5-cleaved permutants.

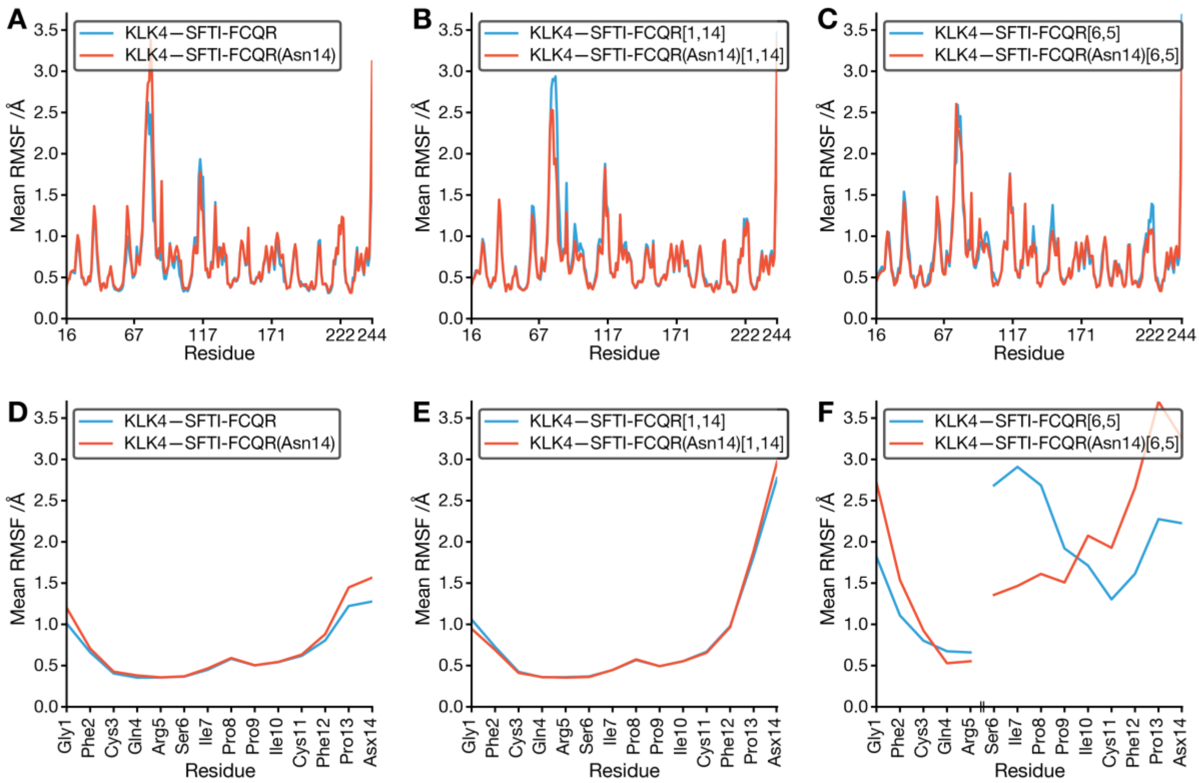


**Supporting Figure 3. Mean Cα-RMSF across multiple runs.** Root mean square fluctuation (RMSF) plots of Cα atoms in simulations with **(A, D)** bicyclic permutants of the SFTI inhibitor, **(B, E)** 1,14-acyclised permutants, and **(C, F)** 6,5-cleaved permutants. Top row contains KLK4 residues, bottom row contains SFTI residues. A zoom-in on the region containing KLK4 residues 186−208 is shown in Supporting Figure 3.

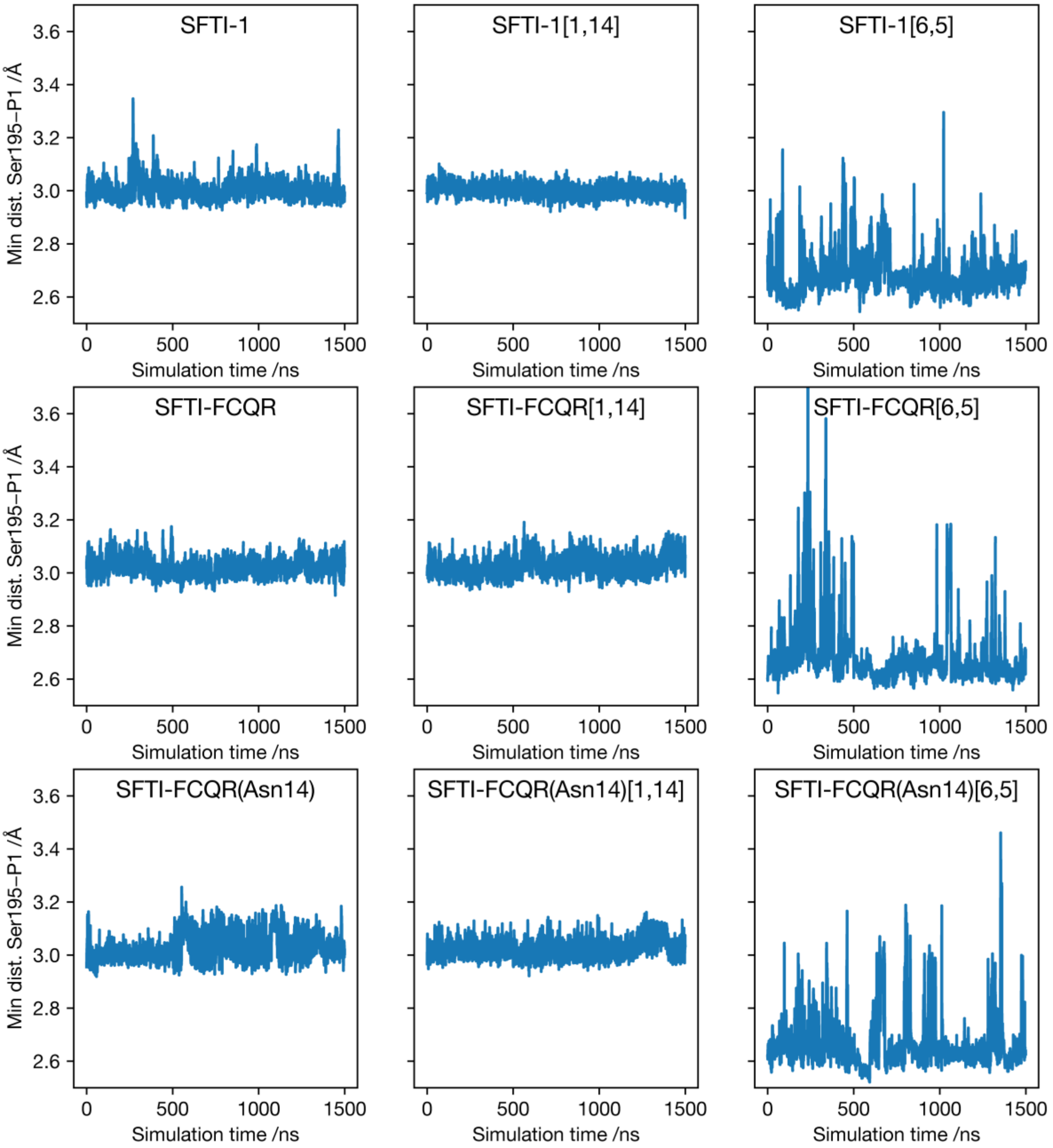


**Supporting Figure 4. Minimum distance between KLK4 Ser195-Oγ and SFTI-derivative P1 residue throughout simulations.** The minimum distance between any atom of the P1 residue sidechain, and the catalytic oxygen of KLK4, displayed as a 1 ns rolling-average. All 500 ns simulations are concatenated to create the plot. Simulations of 6,5-cleaved SFTI permutants showed that P1 residues were closer on average to the catalytic serine, yet more mobile in the S1 pocket than permutants with intact scissile bonds.

**Supplementary Table 1. Intermolecular hydrogen bond distances in KLK4–FCQR(Asn14) and KLK4–FCQR(Asn14)[*1,14*][*6,5*] complexes**, as calculated by PISA. Blank values indicate that a H-bond was not observed.

| **SFTI atom** | **KLK4 atom** | **H-bond distance /Å** | |
|---|---|---|---|
| | | KLK4− SFTI-FCQR(Asn14) | KLK4− SFTI-FCQR(Asn14)[*1,14*][*6,5*] |
| Gly1-O | Ala218-N | 2.99 | 2.97 |
| Cys3-O | Gly216-N | 3.05 | 3.14 |
| Cys3-N | Gly216-O | 2.84 | 2.78 |
| Gln4-Nε2 | Tyr94-Oη | 3.12 | 3.08 |
| Arg5-N | Ser214-O | 3.13 | 3.11 |
| Arg5-Nη1 | Asp189-Oδ1 | 3.06 ** | 2.95 ** |
| Arg5-Nη1 | Ser190-O | 3.15 | 3.11 |
| Arg5-Nη1 | Ser190-Oγ | 2.77 | 2.85 |
| Arg5-Nη1 | Asp189-Oδ2 | 3.40 ** | 3.34 ** |
| Arg5-Nη2 | Lys217-O | 3.13 | 2.96 |
| Arg5-Nη2 | Asp189-Oδ2 | 3.34 ** | 3.32 ** |
| Arg5-O | Gly193-N | 2.73 | 2.81 |
| Arg5-O | Ser195-N | 2.95 | 3.02 |
| Arg5-OXT | His57-Nε2 | | 3.12 |
| Ser6-N | Ser195-Oγ | 3.06 | −; *Ser6 not observed* |
| Ile7-N | Phe41-O | 2.98 | −; *Ile7 not observed* |

**H-bond is considered a salt-bridge.

**Supplementary Table 2. Buried surface area in KLK4−SFTI interface.** Surface areas of SFTI residues buried against KLK4, as calculated by PDBePISA. Values in Å$^2$.

| **SFTI Residue** | **KLK4− SFTI-FCQR(Asn14)** | **KLK4− SFTI-FCQR(Asn14)[*1,14*][*6,5*]** |
|---|---|---|
| Gly1 | 30.25 | 37.58 |
| Phe2 | 134.25 | 120 |
| Cys3 | 47.22 | 47.14 |
| Gln4 | 97.22 | 101.68 |
| Arg5 | 231.86 | 275.23 |
| Ser6 | 51.76 | *Ser6 not observed* |
| Ile7 | 140.68 | *Ile7 not observed* |
| Pro8 | 1.01 | *Pro8 not observed* |
| Pro9 | 24.95 | 6.52 |
| Ile10 | 22.42 | 66.31 |
| Cys11 | 0 | 0.86 |
| Phe12 | 49.94 | 62.58 |
| Pro13 | 0 | 0 |
| Asn14 | 28.66 | 0.5 (*No sidechain observed)* |
